# Polymer-Conjugated Carbon Nanotubes for Biomolecule Loading

**DOI:** 10.1101/2021.07.22.453422

**Authors:** Christopher T. Jackson, Jeffrey W. Wang, Eduardo González-Grandío, Natalie S. Goh, Jaewan Mun, Sejal Krishnan, Markita P. Landry

## Abstract

Nanomaterials have emerged as an invaluable tool for the delivery of biomolecules such as DNA and RNA, with various applications in genetic engineering and post-transcriptional genetic manipulation. Alongside this development, there has been an increasing use of polymer-based techniques, such as polyethyleneimine (PEI), to electrostatically load polynucleotide cargoes onto nanomaterial carriers. However, there remains a need to assess nanomaterial properties, conjugation conditions, and biocompatibility of these nanomaterial-polymer constructs, particularly for use in plant systems. In this work, we develop mechanisms to optimize DNA loading on single-walled carbon nanotubes (SWNTs) with a library of polymer-SWNT constructs and assess DNA loading ability, polydispersity, and both chemical and colloidal stability. Counterintuitively, we demonstrate that polymer hydrolysis from nanomaterial surfaces can occur depending on polymer properties and attachment chemistries, and describe mitigation strategies against construct degradation. Given the growing interest in delivery applications in plant systems, we also assess the toxicity of polymer-based nanomaterials in plants and provide recommendations for future design of nanomaterial-based polynucleotide delivery strategies.

## 1. Introduction

Genetic engineering is a critical component of biomedical research, healthcare, biopharmaceuticals, and agriculture. Central to these applications is the ability to deliver biomolecular cargoes such as DNA, RNA, or proteins, to cells. This delivery challenge affects the efficiency of resulting genetic transformations and the ease and throughput of advancing bioengineering applications. In particular, the low biomolecular cargo delivery efficiencies in plant systems motivates the development of tools for more effective delivery of biomolecular cargoes such as polynucleic acids. Nanomaterial-polymer conjugates have the potential to overcome many of the shortcomings of conventional delivery systems, including low efficiency, species dependence, limited cargo types, and tissue damage.^[1]^

Nanoparticles have been widely used in both mammalian and plant systems for the delivery of biomolecular cargoes. For example, conjugated polymer nanoparticles were shown to effectively penetrate tobacco BY-2 protoplasts within two hours of delivery for small interfering RNA (siRNA) delivery and gene knockdown.^[2]^ Similarly, new polymer compositions for DNA delivery have been demonstrated in moss and tobacco protoplasts, where delivery efficiency is dependent on the chemical structure and molecular weight of the polymer carriers.^[3]^ The formation of ionic complexes that combine a polycation with a cell-penetrating peptide have also enabled the delivery of DNA to intact leaf cells.^[4]^ In both plant and mammalian systems, PEI remains one of the most commonly used cationic polymers for DNA delivery. The delivery capabilities of these PEI-based systems have been broadly attributed to their ability to escape endosomes via a “proton sponge” mechanism. When placed in an acidic endosomal environment, the polymer’s amine groups become increasingly protonated, leading to a buffering effect. As protons (and typically chloride ions, which maintain charge neutrality) enter the vesicle, they cause osmotic swelling and rupture, freeing the nanoparticle and/or its cargo.^[5]^ However, as is the case with many cationic polymers, aggregation of PEI-DNA, which occurs largely due to hydrophobic interactions, limits their utility for gene delivery.^[6]^ Furthermore, the high charge densities present in cationic polymers such as PEI can induce cytotoxicity.^[1,7]^ Mitigating techniques, including cationic polymer cross-linking, chemical modification of the cationic polymer, and modulation of DNA structure can more effectively condense DNA to increase transfection efficiency and limit toxicity.^[8–11]^

Towards these ends, the conjugation of cationic polymers such as PEI to nanoparticles has been demonstrated to improve transfection efficiency, relative to free PEI polymers, in mammalian cells.^[12,13]^ Importantly, particle size and zeta potential absolute magnitude are key for internalization of nanoparticles within an organelle.^[14]^ Early reports have demonstrated the use of Au-PEI nanoparticles to bind RNA via electrostatic interaction and deliver the cargo in mammalian cells with cytocompatibility and improved gene silencing compared to polymer alone.^[13]^ More recent reports have combined low-dimensional nanomaterials, such as SWNTs, with cationic polymer systems for delivery in diverse plant tissues and mammalian cells.^[15,16]^

In spite of the success of polycationic polymers and their nanomaterial conjugates for polynucleotide delivery, remains a lack of consensus on the optimal design of polymer-nanoparticle complexes that maximize nanoparticle stability, delivery efficiency, and biocompatibility.^[17]^ Herein, we explore the use of polymer-conjugated SWNT nanoparticles and the material properties that govern their use in biomolecule delivery. We next optimize polymer-conjugated SWNT nanoparticle biocompatibility for use in plant systems, which remain less studied than their mammalian counterparts and face additional barriers to cellular entry such as the cell wall. To these ends, we synthesized polymer-SWNTs using a library of cationic polymers conjugated with two chemical techniques to assess their relative functional density, dispersibility, and long-term stability. We further investigated the impact of preparation techniques and cationic polymer design in the stability and DNA loading ability of the resulting polymer-SWNT systems. Finally, we assessed plant stress responses to these nanoparticle-polymer conjugates *in vivo* to provide insight into rational polymer-nanoparticle design to optimize DNA loading, construct stability, and minimize toxicity.

## 2. Results & Discussion

### 2.1 Generating polymer-SWNT constructs

We selected a library of cationic polymers (**Table S1**) commonly used for polynucleotide delivery applications, with ranging physicochemical properties including molecular weight, amine density, and structure: linear vs. branched. We also developed two attachment chemistries to covalently link polymers to the SWNT surface: EDC-NHS and triazine chemistries. For EDC-NHS based polymer attachment, commercially available carboxylic acid functionalized SWNTs (COOH-SWNTs) were modified via EDC-NHS chemistry to form a covalent amide bond to the amine groups of the cationic polymers in our library (**Figure 1a**).^[18]^ The attachment of polymer was confirmed by zeta potential measurements, with a notable change from the initial −50.1 mV for COOH-SWNTs to +67.7 mV after conjugation of a cationic polymer such as 25,000 MW branched PEI (BPEI-25k) (**Figure 1c**), with all zeta potential values listed in **Table 1**.

**Figure 1.**
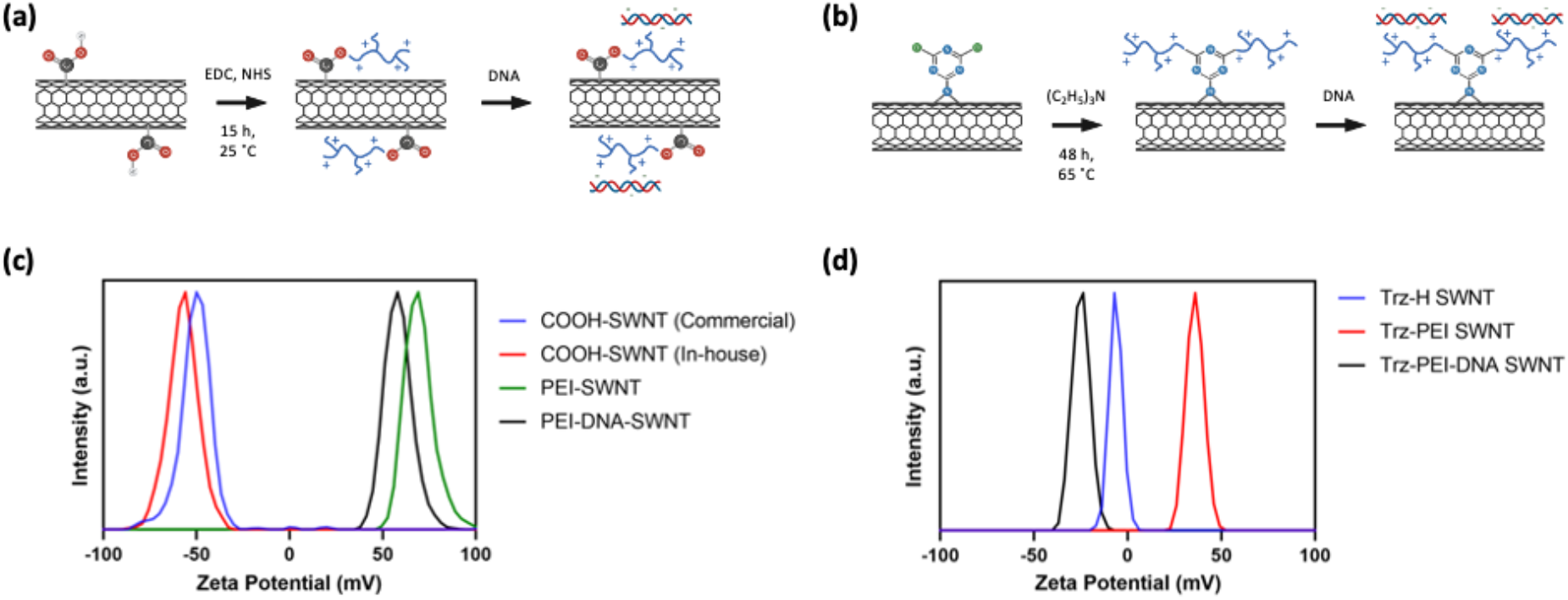
Synthesis and characterization of polymer-SWNTs. (a) Scheme of polymer-SWNT synthesis using EDC-NHS chemistry and subsequent DNA loading. (b) Scheme of polymer-SWNT synthesis using triazine chemistry and subsequent DNA loading. (c) Zeta potential measurements of initial COOH-SWNT constructs, after conjugation with BPEI-25k via EDC-NHS chemistry, and after addition of DNA. (d) Zeta potential measurements of triazine-functionalized SWNTs, after conjugation with BPEI-25k via nucleophilic substitution, and after addition of DNA.

**Table 1.** Zeta potential of polymer-SWNTs.

| **Polymer** | **Polymer-SWNT zeta potential [mV]** |
| --- | --- |
| COOH-SWNT (no polymer) | - 50.1 |
| Branched polyethylenimine (800) | + 54.7 |
| Linear polyethylenimine (500) | + 45.6 |
| Branched polyethylenimine (750k) | + 65.9 |
| Branched polyethylenimine (25k) | + 71.6 |
| Branched polylysine | + 54.0 |
| Linear polyethylenimine (800) | + 59.9 |
| Low hydrophobic modified branched polyethylenimine | + 72.8 |
| Medium hydrophobic modified branched polyethylenimine | + 69.2 |

In the second method, triazine-functionalized SWNTs (Trz-SWNTs) were synthesized from pristine SWNTs via a re-aromatization reaction to generate triazine groups on the SWNT surface.^[19]^ These Trz-SWNTs were further functionalized via a nucleophilic substitution of the chlorine on the triazine with a polymer amine group to create Trz-SWNTs with a covalently attached BPEI-25k polymer (**Figure 1b**). Given the prolific use of this BPEI-25k polymer for polynucleotide delivery applications over others in our library, we only synthesized the BPEI-25k polymer-SWNT complex with this triazine-based chemistry for comparison against EDC-HNS based polymer attachment.The attachment of the BPEI-25k polymer was confirmed by zeta potential measurements, with an increase in zeta potential from −6.30 mV to +36.0 mV after conjugation of BPEI-25k (**Figure 1d**).

### 2.2 Improved functionalization density and removal of amorphous carbon

The efficiency of polymer-SWNT conjugation depends on the purity and the density of COOH functional groups on the COOH-SWNT starting material. Thus, we first implemented thermogravimetric analysis (TGA) of COOH-SWNTs, as received from the supplier, to assess the purity of the COOH-SWNT starting material. Previous literature indicates that both pristine SWNT and COOH-SWNT are thermally stable below 600 °C.^[20,21]^ Upon heating samples to this temperature and accounting for the removal of impurities such as excess solvent below 150 °C, we observe a 81.6% loss in mass. Based on previous literature, we attributed this mass loss to the combustion of amorphous carbon in the sample (**Figure 2a**).^[20,22]^

**Figure 2.**
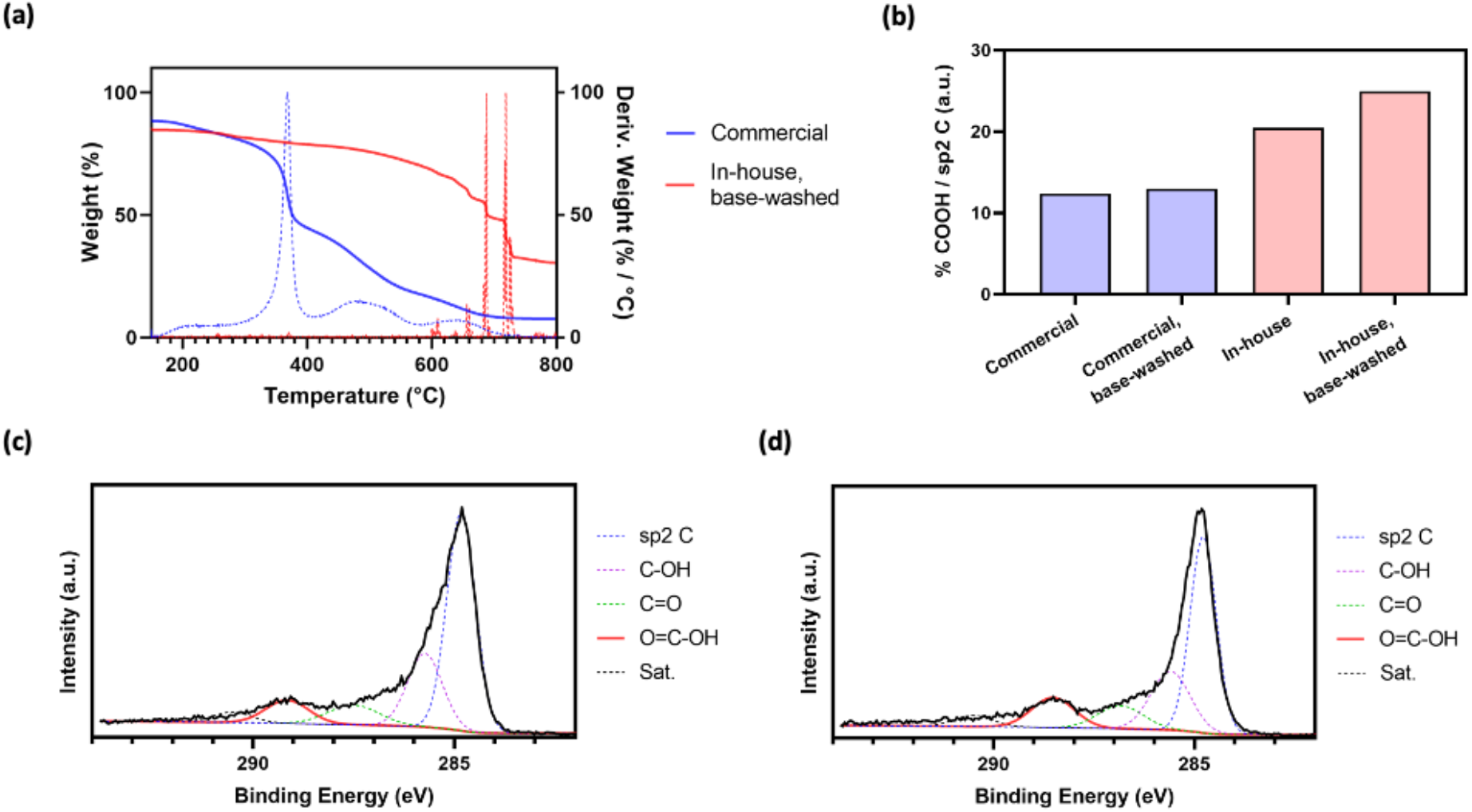
Quantification of amorphous carbon and SWNT carboxylation. (a) TGA measurements of COOH-SWNTs either purchased commercially, carboxylated in-house via reflux in nitric acid, and washed with 1.0 M NaOH. (b) Bar plot displaying carboxyl group peak area, normalized in relation to sp^2^ C peak area in C1s XPS spectra. (c) C1s XPS spectra of commercially purchased COOH-SWNT. (d) C1s XPS spectra of in-house carboxylated SWNTs after washing with 1.0 M NaOH.

To mitigate manufacturer variability, we performed an in-house carboxylation reaction by refluxing pristine SWNTs in concentrated nitric acid.^[23]^ These SWNTs were subsequently washed via vacuum filtration and characterized. The negative zeta potential of our in-house carboxylated SWNTs was −56.9 mV, which is consistent with that of commercially available COOH-SWNTs (−50.1 mV) (Figure 1c) and provides one confirmation of a successful reaction. X-ray photoelectron spectroscopy (XPS) characterization of this product (**Figure S2a**) further confirmed a high degree of in-house produced COOH-SWNT carboxylation, notably higher than that of the commercially-procured COOH-SWNT (**Figure 2b**). COOH-SWNTs synthesized via this technique demonstrated notably less (19.3%) mass loss via TGA analysis, representing a more than four-fold increase in purity compared to commercially purchased SWNTs (**Figure 2a**).

To test whether carboxylation resides predominantly on the SWNT surface compared to on amorphous carbon, we washed commercially purchased carboxylated SWNTs with a 1.0 M NaOH solution to remove amorphous carbon.^[24]^ A colored filtrate was recovered, which has been previously attributed to the presence of oxidation debris.^[25]^ XPS characterization before and after washing suggests that both commercially purchased (**Figure 2c**) and in-house synthesized COOH-SWNTs (**Figure S2a**) contain a high percentage of ester groups (**Table S2**). The subsequent decrease in these ester groups after a base wash treatment suggests that these functional groups are primarily located on amorphous carbon, rather than on the SWNT surface. This result is consistent with previous published literature, which suggests that upon reaction with concentrated acid, oxidation debris from amorphous carbon coats the SWNT walls, preventing covalent functionalization of the SWNT surface.^[25,26]^ As a practical result, it is likely that a majority of carboxyl- functionalized carbon material, which is subsequently conjugated to cationic polymers, is largely removed during wash steps. Any amorphous carbon that is not removed still adsorbs DNA but lacks the material properties, including tensile strength and high aspect ratio, that have been shown to enable DNA delivery.^[15,27,28]^ Importantly, COOH-SWNTs synthesized via an in-house carboxylation reaction followed by base wash demonstrated the highest degree of carboxylation of all treatments tested (**Figure 2b, 2d**), and are thus the best suited starting material for downstream delivery applications.

### 2.3 Removal of unreacted residual polymer

In this study, we tested the conjugation of eight cationic, amine-containing polymers to SWNTs: three branched PEI (BPEI; 800, 25k, and 750k Da), two linear PEI (LPEI; 500 and 800 Da), two hydrophobically modified branched PEI (low-phi-BPEI, low degree of modification, 25-30k Da; and med-phi-BPEI, medium degree of modification, 1,500-2,000 Da), and a branched polylysine (3,500 Da) (**Table S1**) using the aforementioned EDC-NHS chemistry. It has been previously demonstrated that free polymer will bind DNA in solution, preventing its adsorption to nanoparticles of interest.^[6,9–11]^ Furthermore, polymer-DNA constructs have shown limited success for delivery of DNA in whole plant systems due to barriers such as cell membranes or the plant cell wall.^[4]^ Therefore, the removal of unreacted polymer is critical for the viability of polymer-SWNT nanomaterials for DNA delivery. To this end, following the reaction of 1 mg of functionalized COOH-SWNTs with cationic polymers using EDC-NHS chemistry, we tested the efficacy of various washing methods in their ability to remove the large excess of unreacted free polymer and recover pure polymer-SWNT product. First, polymer-SWNT constructs were spin washed via centrifugation at high speed through a 100 kDa spin filter until only 1 mL of solution remained. 4 mL of water was added to the remaining solution and this water wash was repeated a total of six times. The filtrate containing free polymer was collected after each wash step. Alternatively, polymer-SWNT constructs were washed via vacuum filtration through a fritted filter with a 0.45-μm PTFE membrane. An equivalent volume of water to that used during the spin wash process (~4 mL) was used during each wash step, and the filtrate containing free polymer was collected after each step for a total of six times.

To test the purity of the polymer-SWNT samples as a function of the wash step, filtrates from each wash were added to solutions of plasmid DNA and run on an agarose gel. If any free polymer was to be present in the filtrate, it would bind to the plasmid DNA and result in retention of DNA from running into the gel. Indeed, we observe no bands after the first wash, indicating the presence of free polymer that binds the plasmid and prevents its migration into the gel during electrophoresis (**Figure 3a**). By the sixth wash, we no longer observe polymer in the filtrate solution, regardless of polymer type, as indicated by the migration of plasmid through the gel equidistant to that of the control free plasmid (**Figure 3b**). Testing of the filtrate after each wash step for our BPEI-25k polymer-SWNT construct demonstrates that the filtrate is largely free of polymer by the fourth wash step (**Figure 3c**).

**Figure 3.**
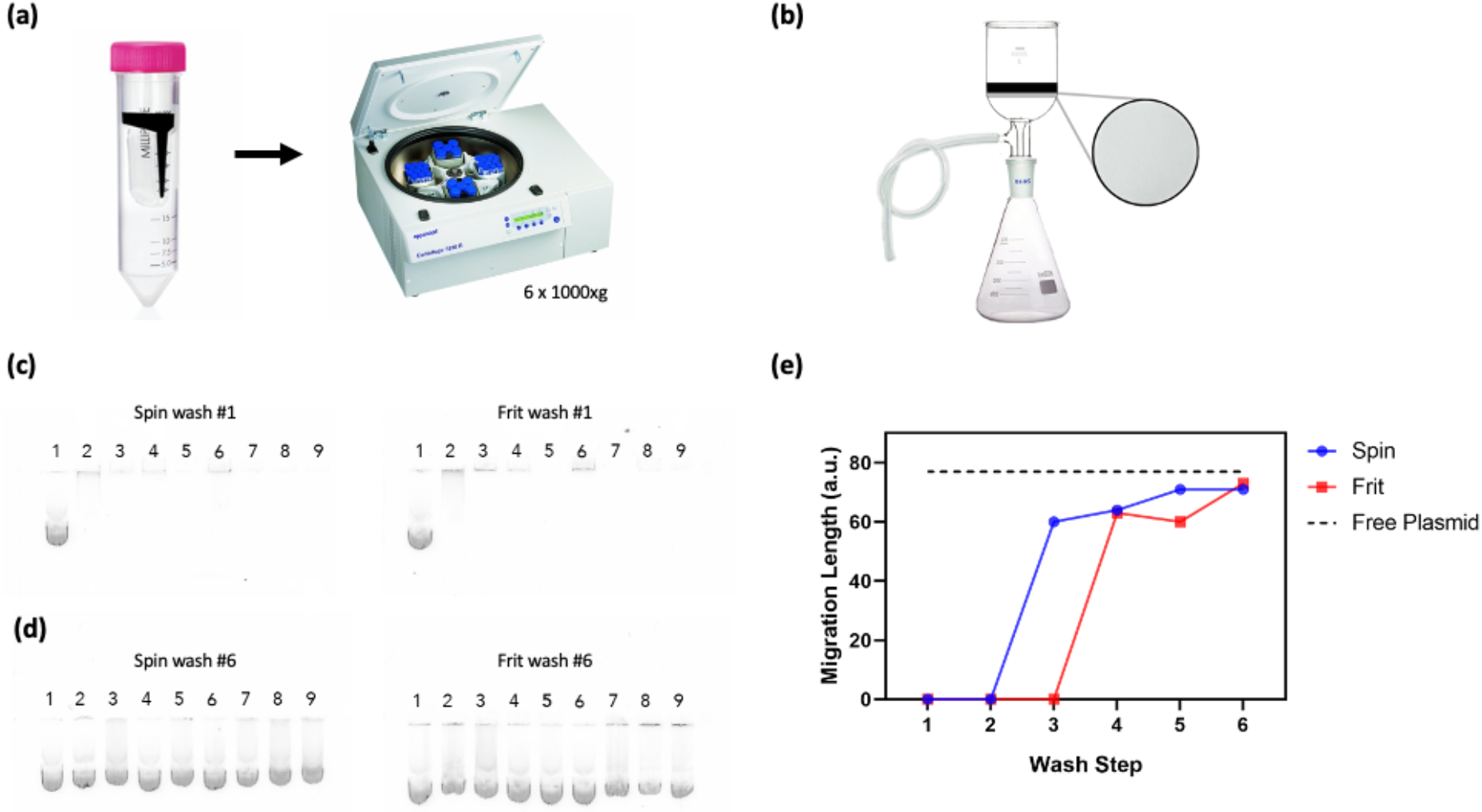
Quantification of free polymer removal from polymer-SWNT complexes. (a) Schematic of polymer-SWNT washing via spin filtration. (b) Schematic of polymer-SWNT washing via vacuum frit filtration. (c) Filtrate from the first polymer-SWNT wash step loaded with DNA and run on an agarose gel for all polymers. (d) Filtrate from the sixth polymer-SWNT wash step loaded with DNA and run on an agarose gel for all polymers. From left to right: (1) free plasmid, (2) LPEI-500, (3) LPEI-800, (4) BPEI-800, (5) low-phi-BPEI, (6) branched polylysine, (7) med-phi-BPEI, (8) BPEI-750k, (9) BPEI-25k. (e) Measurements taken after each wash step for BPEI-25k polymer-SWNTs show a steady increase in the plasmid migration distance, corresponding to a decrease in free polymer, after each wash step.

We further tested the effects of the pH of the wash solution to optimize removal of free polymer (**Figure S4**). In spite of its widespread use, the protonation state of PEI is not well understood; previous studies have suggested that approximately 55% of amine groups are protonated under physiological conditions (pH ~7.4).^[29]^ Wash treatments at pH levels both above and below this pH 7.4 threshold could cause differentially protonated amine groups, impacting the solubility and thus ability of PEI polymers to be removed during the wash process. Generally, we observe that higher pH washes corresponded to a lower zeta potential of the final purified polymer-SWNT sample. We hypothesize that this is due to the poor removal of free polymer, which has a zeta potential that ranges from neutral to weakly positive (**Figure S5**). This hypothesis is further confirmed by the larger size observed by DLS for constructs with the lowest zeta potential, which can also likely be attributed to a large amount of residual polymer in solution and aggregation of the final product.

### 2.4 Probing long-term stability of polymer-SWNTs

Once we confirmed the covalent conjugation of polymers to the SWNT surface and their successful purification from residual unreacted polymer, we examined the long-term stability of polymer-SWNT conjugates in water. Given the challenges of nanoparticle use in diverse biological environments, including biofouling via protein adsorption, loss of colloidal stability, and toxicity, the stability of the nanomaterial polymer bond is critical for a diverse array of applications.^[30]^ For delivery applications, the strength of covalent polymer attachment to nanoparticle surfaces is commonly assumed to to be robust against breakage over conditions relevant to polynucleotide delivery, however, our stability assays below suggest otherwise.

To test the long-term stability of the polymer-SWNT conjugate, we synthesized polymer-SWNT constructs as described above, including 6 water wash steps, to confirm our final product contained purified polymer-SWNTs. Subsequently, we allowed our polymer-SWNT constructs to age in water at ambient conditions for 30 days post-synthesis. As previously demonstrated, we are able to successfully remove all unreacted free polymer after synthesis; therefore, all subsequent measurements can be attributed to polymer that has dissociated from the nanomaterial surface over time. Zeta potential measurements of polymer-SWNT samples conducted at the start and end of this 30 day time period show a substantial decrease in zeta potential, with the emergence of peaks corresponding to less positively charged particles as a function of time (**Figure 4a**). We attribute the appearance of these secondary peaks to both free polymer that is no longer conjugated to our SWNT surface as well as polymer-SWNT conjugates with a decreased amount of attached polymer. We do not observe any peaks at a negative zeta potential that would correspond to SWNTs without any bound polymer (**Figure 1c**), indicating that there is still a substantial amount of polymer attached to our constructs regardless of polymer type. We further confirmed this polymer-SWNT bond instability by XPS analysis, where we we see a notable decrease in the N1s peak in an aged polymer-SWNT sample compared to a freshly synthesized batch (**Figure 4b**), providing further evidence of the loss of polymer from the SWNT surface via hydrolysis of the polymer-SWNT covalent bond over time. Lastly, to further confirm polymer-SWNT degradation over time, a newly synthesized sample of polymer-SWNT was flash-frozen in liquid nitrogen and stored at −80 °C for 30 days. After being thawed, the zeta potential of this sample showed minimal polymer desorption (**Figure 4c**).

**Figure 4.**
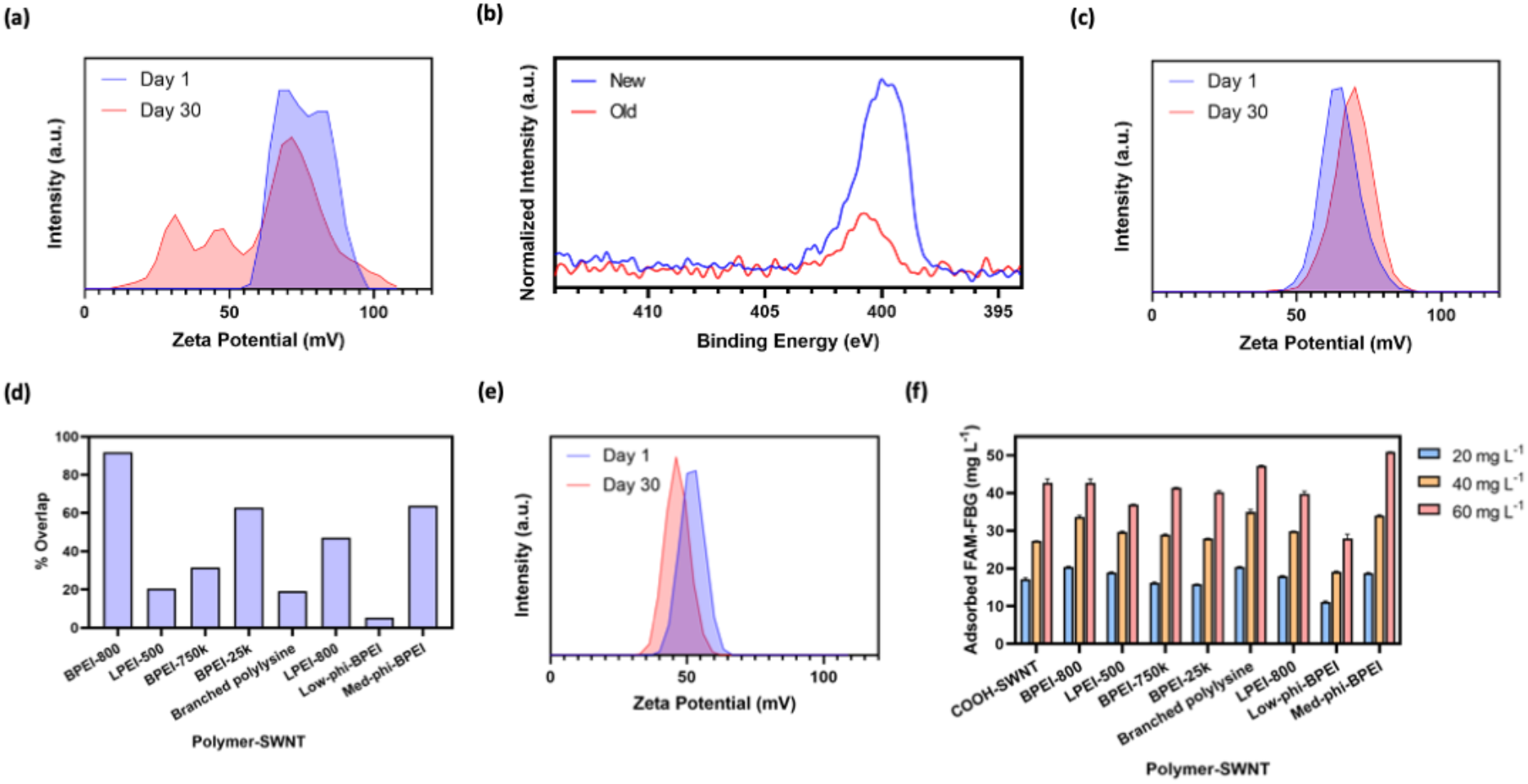
Long-term stability of polymer-SWNT nanoparticles, and protein adsorption to nanoparticles for different polymer attachments. (a) Zeta potential measurements of BPEI-25k polymer-SWNT immediately after synthesis and after 30 days. (b) N1s XPS spectra of a fresh and aged BPEI-25k polymer-SWNT sample, normalized to the respective C1s peak. (c) Zeta potential of a BPEI-25k polymer-SWNT immediately after synthesis and after storage at −80 °C for 30 days. (d) Quantification of the area under the curve overlap in zeta potential spectra peaks between days 1 and 30 for each polymer-SWNT construct. (e) Zeta potential measurements of BPEI-25k-Trz-SWNT immediately after synthesis and after 30 days. (f) Concentration of adsorbed FAM-FBG on 5 μg mL^−1^ polymer-SWNT. Initial concentrations of FAM-FBG added to solution were 20, 40, and 60 μg mL^−1^ respectively. Error bars represent standard deviation of the mean (N = 3).

Each polymer showed different rates of hydrolysis from the SWNT surface (**Figure S6**). To quantify the relative loss in polymer for each polymer-SWNT sample, we calculated the percentage overlap between zeta potential peaks measured at days 1 and 30 for each polymer-SWNT construct (**Figure 4d**), where a higher overlap value represents greater polymer-SWNT stability. We find that both very small and very large PEI polymers disassociate from the SWNT surface rapidly relative to their medium-sized counterparts. In addition, polymers with lower amine density, including LPEI-500, LPEI-800, and branched polylysine, likewise showed poor stability relative to polymers with high amine density, which may be attributed to the relatively lower availability of amines for covalent conjugation to the SWNT surface. We also observe more polymer loss from the SWNT surface for low- versus medium-phi-BPEI polymers, suggesting that increasing polymer hydrophobicity may aid in long-term polymer-nanoparticle stability. These results indicate that we can attribute polymer-SWNT stability to several factors, including polymer amine density (higher is better), sterics (less steric hindrance is better), and polarity (more hydrophobicity is better). Together, our experiments suggest that a compromise between polymer size and structure may be ideal, as exemplified by the BPEI-800 polymer which showed uniquely high stability on the SWNT surface over time. Taken together, based on our experiments and previous literature, we attribute the loss of polymer from the SWNT surface to hydrolysis of the amide bond over time.^[31]^

To probe the effect of alternative bonding chemistry on polymer stability, we performed the same time course study of polymers attached to SWNT with a different attachment chemistry. Specifically, instead of the commonly-used EDC-NHS chemistry, we attached the BPEI-25k-PEI polymer to SWCNT via triazine chemistry. This chemistry proceeds via a covalent functionalization reaction that re-aromatizes SWNT defect sites to restore the original, pristine SWCNT lattice, and yields functional groups on approximately 1.64% of carbons on the SWNT surface.^[19]^ After 30 days at ambient conditions, we observe minimal change in the zeta potential (**Figure 4e**), despite the fact that the EDC-NHS chemistry performed above proceeds with COOH-SWNT starting material containing the same or more functional group density on the SWNT lattice than triazine chemistry. These results suggest that triazine-based attachment chemistries could be more viable for applications where long-term stability of the polymer-nanoparticle construct is necessary.

Lastly, biofouling considerations are often overlooked for polynucleotide delivery applications. Specifically, spontaneous protein adsorption to nanoparticle surfaces can alter the physicochemical properties of the polymer-nanoparticle complex, generating adverse outcomes for successful DNA or RNA loading and delivery. To understand the impact of different covalently linked polymers on protein adsorption to the SWNT surface, we performed an assay to test the adsorption of fibrinogen, a protein known to be highly involved in the formation of nanoparticle coronas.^[32]^ We have previously demonstrated that the fluorescence of FAM-labeled fibrinogen (FAM-FBG) is quenched when this species adsorbs to a SWNT surface.^[33]^ Therefore, we can use this solution-phase and real-time ligand binding assay to quantify the amount of protein that adsorbs onto a nanoparticle surface. When compared to COOH-SWNT across a range of FAM-FBG concentrations, we consistently observe that only one of our polymer-SWNT constructs, the low-phi-BPEI, mitigates against protein adsorption (**Figure 4f**). We hypothesize that this anti-biofouling effect is due to a combination of both the polymer’s hydrophobic modifications and large size, which together may prevent protein from binding to the SWNT surface.^[30]^

### 2.5 DNA loading on polymer-SWNT constructs

We next investigated the ability of our polymer-SWNT nanomaterials to electrostatically bind plasmid DNA. Measurements taken before and after addition of DNA show a decrease in zeta potential after the addition of DNA, as expected due to the negative charge of DNA (**Figure 5a**). For the lowest molecular weight polymer-SWNT construct (LPEI-500-SWNT), we observe a negative final zeta potential for the DNA-polymer-SWNT mixture. This decrease in zeta potential, which was also observed to a lesser extent for BPEI-800-SWNT and branched polylysine-SWNT following DNA addition, is also accompanied by increase in size as measured by DLS, which is attributed to aggregation of these nanoparticles (**Figure 5b**). We hypothesize that the lower molecular weight of these PEI polymers results in less polymer mass available per conjugation site on the SWNT surface, and as a result, a decreased ability to bind DNA as effectively as larger polymers.^[11]^ Similarly, the lower density of amine groups in polylysine, especially as compared to BPEI, likely inhibits its ability to bind to the SWNT surface through EDC-NHS chemistry.

**Figure 5.**
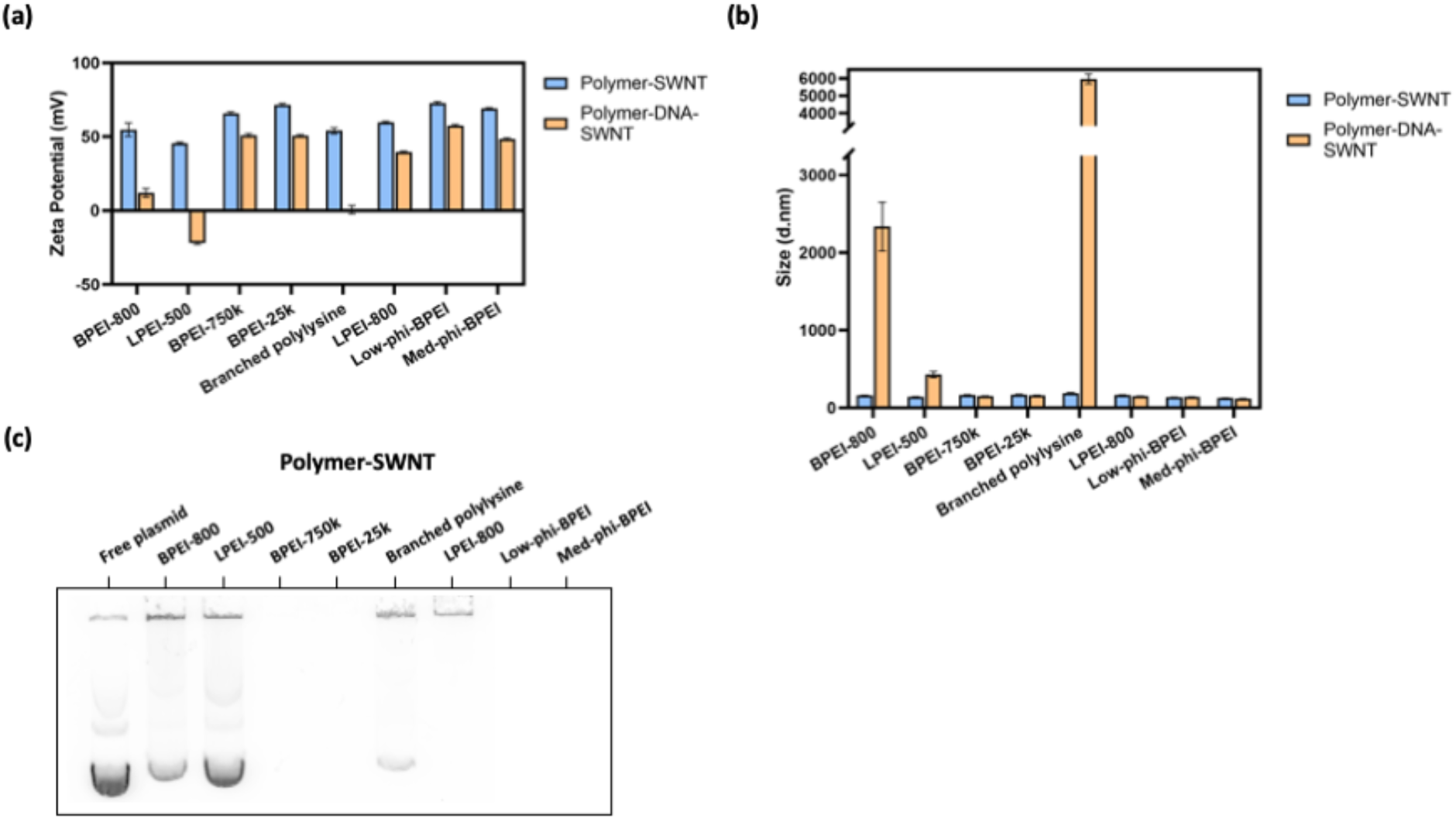
DNA loading capacity of polymer-SWNTs. (a) Zeta potential measurements, in water, of polymer-SWNT constructs, before and after addition of DNA. (b) DLS measurements, in water, of polymer-SWNT constructs, before and after addition of DNA. Error bars represent standard deviation of the mean (N = 3). (c) Agarose gel of polymer-SWNTs (10 μg mL^−1^) loaded with DNA (10 μg mL^−1^).

To better understand the relative loading ability of these various polymer-SWNT conjugates, we loaded DNA on polymer-SWNTs at a 1:1 mass ratio, and next loaded DNA-polymer-SWNT samples in an agarose gel (**Figure 5c**). Due to the size and net charge of DNA successfully loaded on polymer-SWNTs, we anticipated that successfully-bound DNA would exhibit retention in the loading well and would not run into the agarose gel during electrophoresis. For several polymer-SWNT constructs (BPEI-750k, BPEI-25k, low-phi-BPEI, and med-phi-BPEI), we do not observe any DNA running into the gel, suggesting these polymer-SWNT effectively load DNA at a 1:1 mass ratio. In contrast, we observed that the DNA loaded on polymer-SWNTs that previously showed a low or negative zeta potential, and colloidal instability, after DNA loading (BPEI-800, LPEI-500, and branched polylysine) ran into the gel during electrophoresis, indicating the presence of free plasmid. These results confirm a large range of variability in the effectiveness of different polymer-SWNT constructs for loading polynucleotides such as plasmid DNA.

### 2.6 Plant stress response upon infiltration with polymer-SWNTs

As previously discussed, cationic polymers such as PEI are known to be toxic in mammalian cells, which severely limits their use in gene delivery.^[6,9,34]^ However, polymer toxicity in plants is less well understood, particularly when used in conjunction with nanoparticle systems. To test the biocompatibility of polymer-SWNTs in plants, we abaxially-infiltrated 50 mg L^−1^of polymer-SWNT nanoparticles into leaves of 5-week old mature *Nicotiana benthamiana (Nb)* plants, a common model laboratory plant species.^[35]^

To assess toxicity, we infiltrated *Nb* leaves with polymer-SWNT nanoparticles and compared differential expression of stress genes in these leaves, relative to leaves infiltrated with COOH-SWNTs. By performing this comparison, we sought to isolate the toxic effect of each polymer-SWNT conjugate relative to the COOH-SWNT starting material. Two days post-infiltration, we harvested leaf tissue and performed qPCR analysis of *pathogenesis-related gene 1* (*PR1A*) upregulation, a known stress gene in *Nb* plants.^[36]^ Quantification of *PR1A* expression shows that areas infiltrated with SWNT-branched PEI polymers exhibit large upregulation of *PR1A* two days after infiltration (**Figure 6c**). This stress response was observed most strongly in leaf tissues infiltrated with higher molecular weight polymer-SWNT conjugates. In contrast, low molecular weight and linear polymer-SWNT conjugates exhibited a relatively low stress response. Interestingly, a low degree of hydrophobic modification for branched PEI polymer-SWNT conjugates significantly reduced upregulation of *PR1A*, which also reduced nonspecific protein adsorption (**Figure 4f**). We hypothesize that a combination of hydrophobic modifications and steric effects from large molecular weight PEI polymer limits the adsorption of proteins in plant media, which in turn mitigates toxicity of the nanoparticle-polymer conjugates. Similar trends have been shown in previous literature, whereby low enhancements in hydrophobicity increase transfection efficiency of PEI polymers.^[11]^ These findings were further confirmed by testing the response of *arabinogalactan protein 41* (NbAGP41), for which the orthologous gene in *Arabidopsis thaliana* (AT5G24105) has previously shown to be downregulated during plant stress response.^[37]^ We found that the low-phi-BPEI and branched polylysine both led to upregulation of this gene, suggesting their biocompatibility in *Nb* plants (**Figure 6d**). We performed the same tests in *Arabidopsis thaliana* plants, where we observed no notable difference in stress response trends between leaves infiltrated with BPEI-25k-SWNT and DNA-BPEI-25k-SWNT constructs (**Figure S8**).

**Figure 6.**
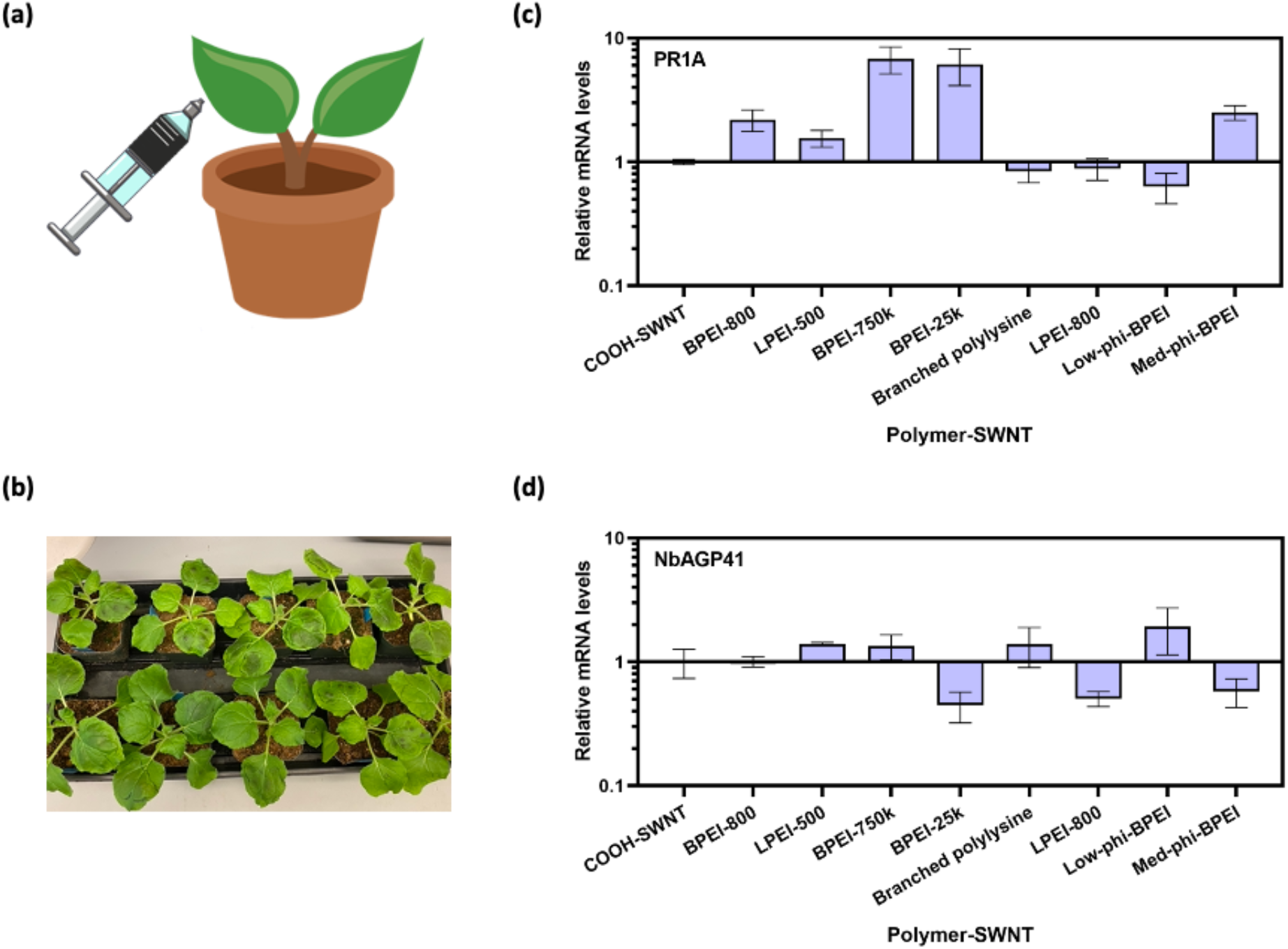
Plant toxicity induced by polymer-SWNT nanoparticles measured by stress gene response in *Nicotiana benthamiana*. (a) Graphic illustration of leaf infiltration with polymer-SWNTs. (b) Plant leaves immediately post-infiltration. (c) qPCR analysis quantifying mRNA fold-change for PR1A gene two days after infiltration. d) qPCR analysis quantifying mRNA fold change for NbAGP41 gene two days after infiltration. Error bars represent standard deviation of the mean (N = 4).

Interestingly, we observe that the low-phi-BPEI-SWNT, which previously demonstrated the lowest levels of protein adsorption, also showed the highest level of biocompatibility as assessed via qPCR. This correlation suggests that protein adsorption to nanoparticles may play an important role in plant stress response, creating an opportunity for the tailoring of polymer properties to enhance or mitigate these effects.

## 3. Conclusion

Despite the widespread use of polymer-nanoparticle conjugates for the delivery of biomolecular cargo, there lacks consensus on what nanocarrier properties maximize their loading ability, polydispersibility, stability, and biocompatibility. To address these issues, we generated and characterized the properties of a library of polymer-SWNT nanoparticles for DNA loading. We found that commercially available carboxylated SWNTs contain a high degree of amorphous carbon, which is detrimental to downstream chemical modification and successful recovery of polymer-SWNT complexes. Therefore, we identified synthetic techniques, including carboxylation via acid reflux and subsequent washing with a basic solution, to aid in removal of amorphous carbon and improve functionalization density on the SWNT surface. Subsequently, we identify that the presence of free polymer, whether residual after covalent conjugation or as a product of hydrolysis from the nanoparticle surface over time, can also inhibit the electrostatic adsorption of polynucleotides to the nanoparticle. We demonstrate successful removal of free polymer and techniques to minimize hydrolysis through polymer selection and storage conditions. By testing different cationic polymers, we demonstrate their differing abilities to load DNA, largely as a function of the polymer size and structure. These material properties also play a role in biomolecule adsorption and plant toxicity, suggesting the need for improved polymer design that can address both of these challenges.

This study further highlights the wide tunability of polymer-SWNT chemistry that can allow for improved biomolecule loading and stability. Our results show successful covalent attachment of a variety of cationic polymers and provide insight into rational polymer design for improved conjugation and electrostatic adsorption of DNA. These findings demonstrate the possibility of new chemistries and material design that can build upon the inherent advantages of nanomaterials such as SWNTs for cargo delivery in biological systems.

## 4. Experimental Section / Methods

### i. Materials

All chemicals unless otherwise noted were purchased from Sigma Aldrich. Branched PEI polymers were purchased from Sigma Aldrich; all other polymers were provided by BASF. Carboxylated SWNTs were purchased from Sigma Aldrich. Raw high pressure carbon monoxide (HiPCO) synthesized SWNTs were purchased from NanoIntegris.

### ii. Synthesis of COOH-SWNTs

Synthesis of COOH-SWNTs was adapted from previous literature.^[23]^ SWNT (20 mg) was combined in a round-bottom flask with 2.6 M HNO_3_ (40 mL). This was connected to a reflux condenser and stirred with a magnetic stirring bar at 120 °C for 12 hours. After being cooled to room temperature, the product was isolated via vacuum filtration and washed sequentially with water, methanol, DMF, NaOH, and water. The solid was lyophilized for storage.

### iii. Synthesis of EDC-NHS polymer SWNTs

Synthesis of EDC-NHS polymer SWNTs was adapted from previously published work.^[15,18]^ COOH-SWNTs were added to water in a 1 mg:1 mL ratio and dispersed via bath (10 min) and probe-tip (30 min, ~30-40 W) sonication. The resulting solution was centrifuged at 18000 xg for 1 h. Afterwards, the supernatant was collected, and the concentration was measured via absorbance at 632 nm with an extinction coefficient of 0.036.

COOH-SWNT (1 mg) was dispersed in 100 mM MES buffer and adjusted to a pH of 4.5 - 6. N- (3-dimethylaminopropyl)-Nʹ-ethylcarbodiimide hydrochloride (EDC) (5 mg) and N-hydroxysulfosuccinimide sodium salt (NHS) (5 mg) were dissolved in 100 mM MES solution (2.5 mL) and added dropwise to the SWNT mixture while stirring. The solution was bath sonicated for 15 minutes and then placed on an orbital shaker at 100 rpm for 45-60 minutes. The product was then washed three times with 0.1X PBS via spin filtration at 300xg for ~8 min through a 100K MWCO filter. Each polymer (20 mg) was dissolved in 0.1X PBS and adjusted to a pH between 7.4-7.6. The SWNT solution was added dropwise to the polymer solution while stirring. The pH was adjusted to a range of 7-8 and the solution was placed on an orbital shaker at 180 rpm overnight.

The resulting product was redispersed via probe-tip sonication (if significantly aggregated) and washed six times with water via spin filtration at 1000xg through a 100K MWCO filter (1-20 minutes each, depending on the polymer). The product was resuspended via bath and probe-tip sonication, centrifuged, and the supernatant was collected. The SWNT concentration was measured via absorbance at 632 nm with an extinction coefficient of 0.036.

### iv. Synthesis of triazine polymer SWNTs

Synthesis of triazine polymer SWNTs with high labeling density (Trz-H) was adapted from previously published work.^[19]^ Trz-H SWNTs (10 mg) were dispersed in dimethylformamide (DMF) (5 mL) and bath sonicated for 15 min. Next, polymer (13.3 mg) and a 1.5 M excess of triethylamine were added and the mixture was stirred at 65 °C for 2 days. The product was washed via centrifugation and re-dispersion in DMF and water (4 mL, two times each). The product was then resuspended in water and washed with water via spin filtration through a 100K MWCO filter six times at 1000xg. The product was resuspended in water and lyophilized for storage.

### v. DNA loading

Plasmid DNA was added to SWNTs at a 1:1 ratio and allowed to incubate at room temperature for 30 min. For plant infiltrations, solutions were diluted with MES delivery buffer to a final volume of 100 μL, with 500 ng each of DNA and SWNT respectively.^[18]^

### vi. DLS and zeta potential measurements

DLS and zeta potential measurements were taken on a Zetasizer Nano ZS (Malvern Instruments). SWNT solutions (with and without DNA) were diluted in water to a concentration of 5 mg L^−1^. Three replicates of at least 20 measurements were obtained for each sample after 2 min equilibration.

### vii. X-ray photoelectron spectroscopy (XPS)

Samples were drop cast onto the surface of a clean silicon wafer. XPS spectra were collected with a PHI 5600/ESCA system equipped with a monochromatic Al Kα radiation source (hν = 1486.6 eV). High-resolution XPS spectra were deconvoluted with MultiPak software (Physical Electronics) by centering the C–C peak to 284.8 eV, constraining peak centers to ± 0.2 eV the peak positions reported in previous literature, constraining full width at half maxima ≤1.5 eV, and applying Gaussian–Lorentzian curve fits with the Shirley background.

### viii. Thermogravimetric analysis (TGA)

TGA measurements were conducted on a TGA 29950 Thermogravimetric Analyzer (TA Instruments). Samples were transferred to an alumina holder and placed in an inert nitrogen atmosphere. The temperature was increased from room temperature to 150 °C, held for 3 hours, then gradually raised to 800 °C before being rapidly cooled. Measurements were taken every two seconds over the course of 17.5 hours. Mass percentage loss was calculated as the difference between measurements at 308 minutes (150 °C) and 758 minutes (600 °C).

### ix. Fluorescence Tracking of Protein Adsorption

FAM fluorophore was conjugated to fibrinogen (FBG) using N-hydroxysuccinimide (NHS) ester chemistry according to previously published work.^[33]^ SWNT and FAM-FBG were mixed in a 1:1 volume ratio, 50 μL total in a 96-well PCR plate (Bio-Rad) and placed in a CFX96 Real-Time PCR System (Bio-Rad). Final concentrations were 5 μg mL^−1^ SWCNT and 20, 40, or 60 μg mL^−1^ FAM-FBG. Scans were collected at the FAM fluorescence channel at 30 s intervals at 22.5 °C. A FAM-FBG fluorescence standard curve was used to convert fluorescence readings to unbound FAM-FBG concentrations.

### x. Gel analysis

For experiments to remove free polymer (**Figure 3a-c**), filtrate (1 μL) was added to DNA (100 ng in 5 μL). Samples were loaded with 6x non-SDS containing loading dye and run in 0.8% agarose at 80 V for 45 min.

For experiments to test the DNA loading ability of our materials (**Figure 5c**), polymer-SWNT (100 ng) was added to DNA (100 ng) and diluted to a total volume of 10 μL. Samples were loaded with 6x non-SDS containing loading dye and run in 0.8% agarose at 80 V for 45 min.

### xi. Plant toxicity measurements

Healthy (5 week old) *N. benthamiana* plants were selected for experiments. For each polymer, four replicates were performed on a single plant. 50 mg L^−1^ polymer-SWNT (100 μL) was injected into plant leaves via needle-less syringe infiltration (**Figure 6a-b**).^[18]^ After 2 days, the leaf tissue was collected. RNA was extracted via a TRIzol reagent and subsequently used for cDNA synthesis and qPCR measurements.

## Acknowledgements

M.P.L acknowledges the support of a Burroughs Wellcome Fund Career Award at the Scientific Interface (CASI), a Stanley Fahn PDF Junior Faculty Grant with Award # PF-JFA-1760, a Beckman Foundation Young Investigator Award, a USDA AFRI award, a USDA NIFA award, and a Foundation for Food and Agriculture Research (FFAR) New Innovator Award. M.P.L. is a Chan Zuckerberg Biohub investigator and an Innovative Genomics Institute Investigator. C.T.J. and J.W.W. acknowledge the support of the National Science Foundation Graduate Research Fellowship. N.S.G. acknowledges the support of the FFAR Fellowship. The authors would like to thank Dr. Ian McFarlane for help analyzing zeta potential spectral overlap. We thank Abraham Martinez and Dr. Alexander Katz for help performing TGA and the Kuriyan Lab for the use of their lyophilizer. We thank BASF for providing most of the polymers used. We acknowledge the support of the Berkeley Nanotechnology Center for XPS. We acknowledge the support of Genomic Science Program grant no. DE-SC0020366 from the DOE Department of Biological and Environmental Research. Figures created with BioRender.com.

## Notes

### Competing Interest Statement

The authors have declared no competing interest.

