## Supplementary material for "Polymer-Conjugated Carbon Nanotubes for Biomolecule Loading": SI

**Table S1.** Polymer characteristics.

| Name | Abbreviation | Molecular weight (Da) |
|---|---|---|
| Branched polyethylenimine (800) | BPEI-800 | 800 |
| Linear polyethylenimine (500) | LPEI-500 | 500 |
| Branched polyethylenimine (750k) | BPEI-750k | 750,000 |
| Branched polyethylenimine (25k) | BPEI-25k | 25,000 |
| Branched polylysine | - | 3,500 |
| Linear polyethylenimine (800) | LPEI-800 | 800 |
| Low hydrophobic modified branched polyethylenimine | low-phi-BPEI | 25,000 - 30,000 |
| Medium hydrophobic modified branched polyethylenimine | med-phi-BPEI | 1,500 - 2,000 |

**Table S2.** Peak deconvolution of XPS spectra of COOH-SWNT. Unit = % Peak Area.

| Peak | Pristine | Commercial | Commercial, base-washed | In-house | In-house, base-washed |
|---|---|---|---|---|---|
| **$sp^2$ C** | 71.37 | 57.01 | 64.09 | 51.21 | 50.34 |
| **C-OH** | 10.80 | 23.16 | 14.52 | 23.99 | 22.60 |
| **C=O** | 7.61 | 9.17 | 8.07 | 11.91 | 10.33 |
| **O=C-OH** | 3.92 | 7.06 | 8.30 | 10.48 | 12.57 |
| **Sat.** | 6.30 | 3.61 | 5.02 | 2.40 | 4.16 |

**Table S3.** Size of nanomaterial constructs, characterized by dynamic light scattering (DLS).

| Functionalized nanomaterial | Size (d.nm) |
|---|---|
| COOH-SWNT (Commercial) | 147.9 |
| COOH-SWNT (in-house) | 237.0 |
| PEI-SWNT | 160.8 |
| PEI-SWNT-DNA | 130.3 |
| Trz-H SWNT | 300.3 |
| Trz-PEI SWNT | 394.0 |
| Trz-PEI-SWNT-DNA | 779.8 |

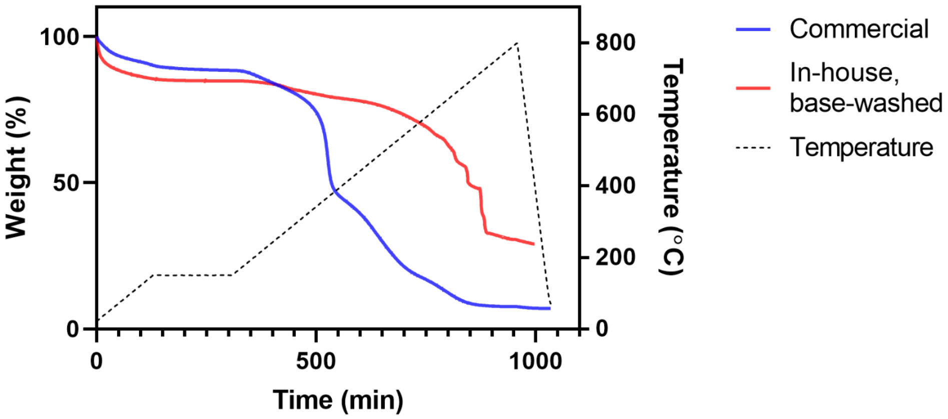


**Figure S1. Thermogravimetric analysis heating profile.** TGA temperature was increased from room temperature to 150 ˚C, held for 3 hours, then gradually raised to 800 ˚C before being rapidly cooled.

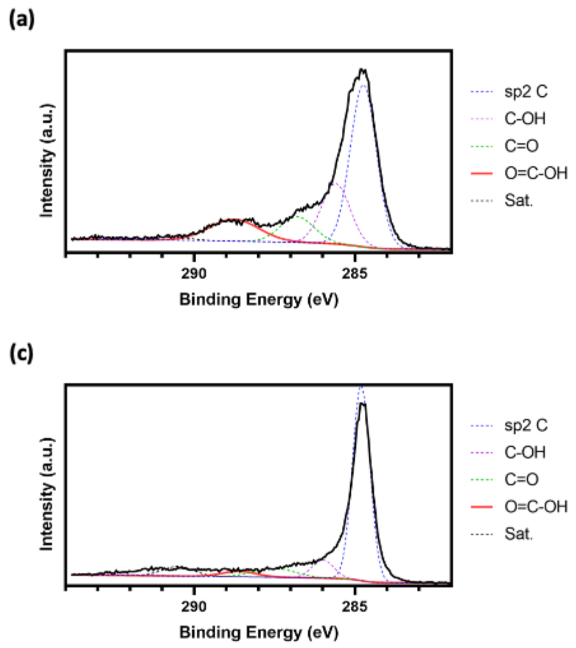


**Figure S2. XPS of carboxylated-SWNT preparations.** (a) XPS C1s spectra of in-house carboxylated COOH-SWNT. (b) XPS C1s spectra of commercially purchased COOH-SWNT after wash treatment with 1.0 M NaOH. (c) XPS C1s spectra of pristine SWNT.

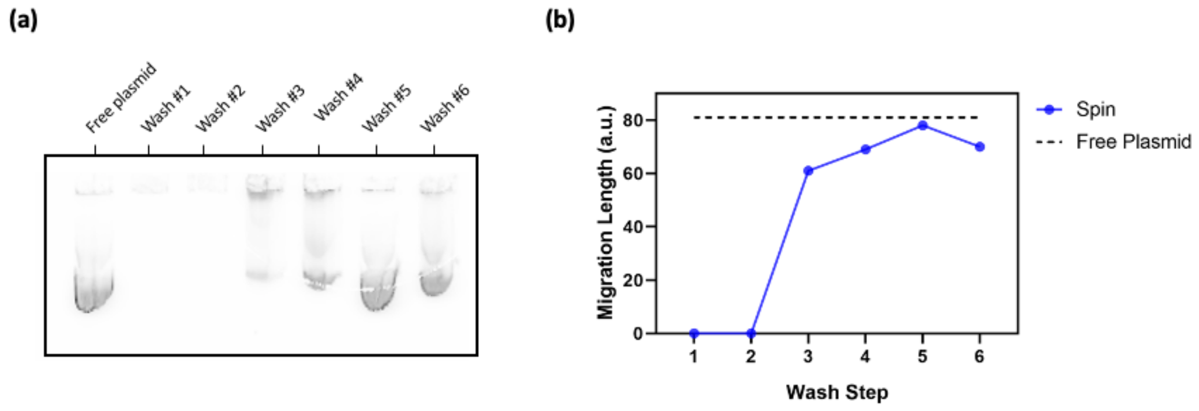


**Figure S3**. **Washing of BPEI-25k polymer.** Free BPEI-25k polymer suspended in water at 8 mg/L was washed six times with water through a 100 kDa spin filter at 1000 x g. The filtrate was collected after each wash step, loaded with DNA, and run on an agarose gel. By the fourth wash step, the filtrate loaded with DNA ran equidistant to free plasmid, suggesting that no free polymer remains in solution.

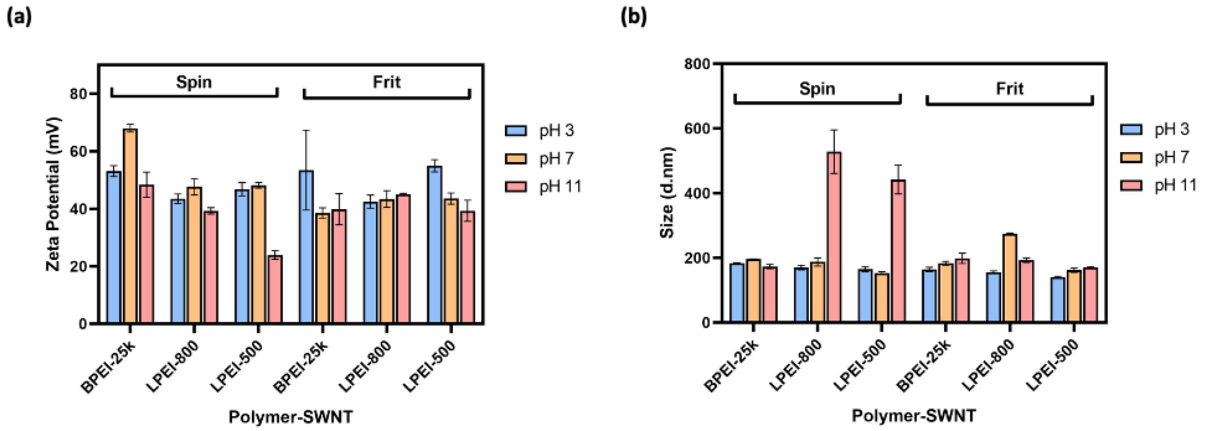


**Figure S4**. **Zeta potential of polymer-SWNTs washed at varying pH.** We attribute a lower zeta potential value to the poor removal of free polymer. (a) Zeta potential of washed polymer-SWNTs. (b) DLS size measurements for washed polymer-SWNTs. Error bars represent standard deviation of the mean (N = 3).

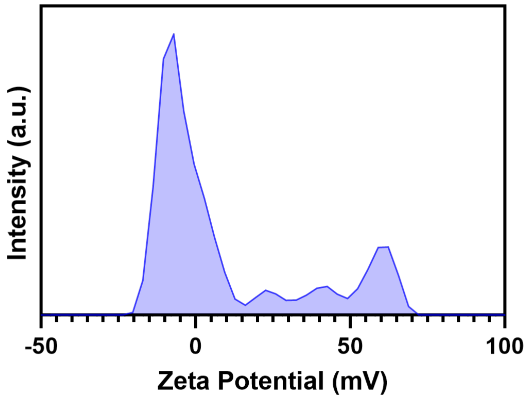


**Figure S5. Zeta potential of BPEI-25k polymer in water.** Free polymer suspended in solution exhibits a neutral or weakly positive charge, likely due to intermolecular interactions. When free polymer is suspended with positively charged polymer-SWNT nanoparticles, this can increase the ionic strength of the suspension and lower the measured zeta potential of the whole solution.

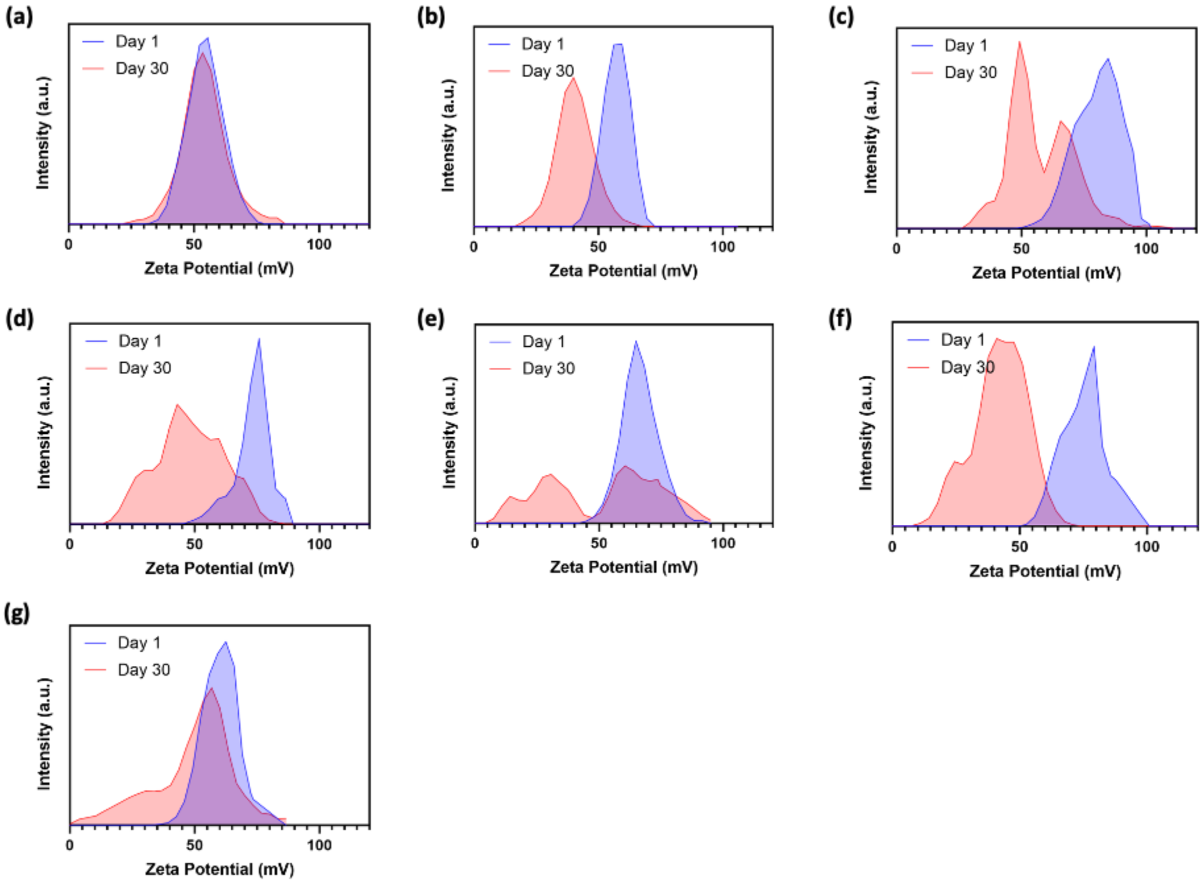


**Figure S6**. **Desorption of polymer from polymer-SWNT nanoparticles over time.** (a) BPEI-800, (b) LPEI-500, (c) BPEI-750k, (d) Branched polylysine, (e) LPEI-800, (f) low-phi-BPEI, (g) med-phi-BPEI.

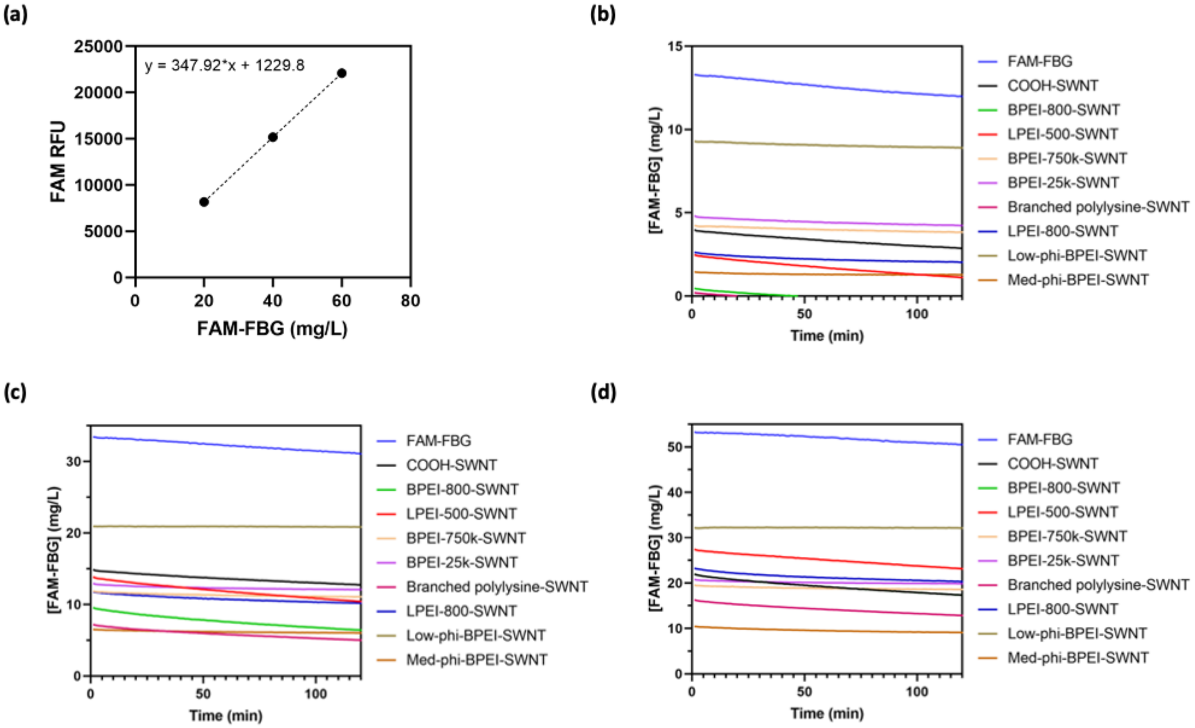


**Figure S7. Quantification of real-time FAM-fibrinogen adsorption to polymer-SWNTs.** (a) FAM-FBG fluorescence calibration curve. (b) Adsorption of 20 µg/mL FAM-FBG determined by quenching of conjugated FAM fluorophore to 5 µg/mL polymer-SWNTs. (c) Adsorption of 40 µg/mL FAM-FBG determined by quenching of conjugated FAM fluorophore to 5 µg/mL polymer-SWNTs. (d) Adsorption of 60 µg/mL FAM-FBG determined by quenching of conjugated FAM fluorophore to 5 µg/mL polymer-SWNTs. The decrease in the FAM-FBG concentration over time across all concentrations can be attributed to self-quenching.

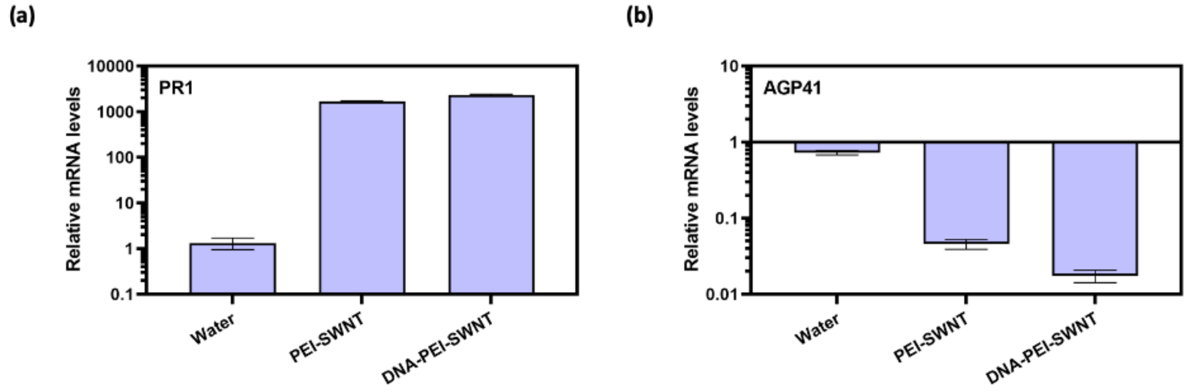


**Figure S8. Toxicity impact of DNA loading on polymer-SWNTs**. We observe no notable difference in trends of mRNA fold change of *arabinogalactan protein 41* (AGP41) between *Arabidopsis thaliana* leaves infiltrated with BPEI-25k-SWNT and DNA-BPEI-25k-SWNT two days post-infiltration. Error bars represent standard error of the mean (N = 3).
